# Rapid metagenomic characterization of a case of imported COVID-19 in Cambodia

**DOI:** 10.1101/2020.03.02.968818

**Authors:** Jessica E. Manning, Jennifer A. Bohl, Sreyngim Lay, Sophana Chea, Ly Sovann, Yi Sengdoeurn, Seng Heng, Chan Vuthy, Katrina Kalantar, Vida Ahyong, Michelle Tan, Jonathan Sheu, Cristina M. Tato, Joseph L. DeRisi, Laurence Baril, Veasna Duong, Philippe Dussart, Erik A. Karlsson

## Abstract

Rapid production and publication of pathogen genome sequences during emerging disease outbreaks provide crucial public health information. In resource-limited settings, especially near an outbreak epicenter, conventional deep sequencing or bioinformatics are often challenging. Here we successfully used metagenomic next generation sequencing on an iSeq100 Illumina platform paired with an open-source bioinformatics pipeline to quickly characterize Cambodia’s first case of COVID-2019.

## Introduction

The ongoing outbreak of a novel coronavirus in China is now evident in 37 other countries as others employ vigilant screening procedures.^1^ As of 24 Feb 2020, 119 SARS-CoV-2 genome sequences from 15 countries are publicly available from over 79,331 confirmed cases.^1,2^ These viral genomes differ by only 0 to 5 mutations, likely representing a single introduction in the human population from an unknown reservoir or intermediate host, potentially bats.^3^ Subsequently, the virus rapidly spread since December 2019. Currently, incubation period and transmissibility are questions of global concern, but difficult to answer given nonspecific and mild symptoms that could represent a variety of respiratory viruses.^4,5^

Genetic characterization of the virus from geographically diverse patient samples is key to infer the rate of spread. Additionally, rapid and complete results from cases worldwide are important for sequence-dependent countermeasures and accurate diagnostics. However, as the virus reaches resource-limited settings in proximity to the outbreak’s epicenter, such as Cambodia, Laos, and Myanmar, there are often logistical challenges in surveillance, contact tracing, sample collection, and biospecimen transport. Information delay is also compounded by a dearth of sequencing and/or bioinformatics expertise in-country, further postponing analysis and dissemination of pathogen genomic information.

Here, in a rapidly implemented response to the novel coronavirus outbreak, we used an unbiased metagenomics next generation sequencing (mNGS) approach on the index COVID-19 case in Cambodia as a tactical demonstration of an efficient workflow to identify and characterize the novel virus. While unbiased sequencing successfully achieved an unambiguous identification of the virus in the absence of any sequence enrichment, complete coverage of the SARS-CoV-2 genome was subsequently achieved by augmenting the mNGS library preparation via a spiked primer enrichment approach.^6^

## Methods

### Viral diagnostic methods

Nasopharyngeal and oropharyngeal swabs (combined into one tube) were collected from a symptomatic patient meeting case definition for possible infection with COVID-19 pneumonia on January 26, 2020. Extraction of viral nucleic acids from clinical sample was performed with a QIAamp Viral RNA Mini Kit (Qiagen #52906) as described by manufacturer. Extracted RNA samples were DNAse-treated and tested via real-time polymerase chain reaction for SARS-CoV-2 using both the Charité Virology and Hong Kong University protocols and confirmed positive on January 27^th^, 2020.^7^

### mNGS preparation and analysis

On January 29, samples were received for cDNA library preparation. Since clinical specimens are obtained from suspected COVID-19 cases as part of the national outbreak response, requirement for informed consent was waived for pathogen identification and characterization. mNGS libraries were prepared from the viral RNA extracted from the sample. cDNA was converted to Illumina libraries using the NEBNext Ultra II DNA Library Prep Kit (E7645) according to the manufacturer’s recommendations. Library size and concentration were determined using the 4150 Tapestation system (Agilent, MA, USA). External RNA Controls 103 Consortium collection (ThermoFisher, 4456740) spike-in controls were used as markers of potential library preparation errors and for input RNA mass calculation. Sample was sequenced on an Illumina iSeq100 instrument using 150 nucleotide paired-end sequencing. A water (“no-template”) control was included in library preparation. Microbial sequences from the sample are located in the National Center for Biotechnology Information (NCBI) Sequence Read Archive.

Raw fastq files were uploaded to the IDseq portal, a cloud-based, open-source bioinformatics platform to identify microbes from metagenomic data (https://idseq.net). The filter steps and microbial data are accessible at http://public.idseq.net/covid-19?utm_source=bioarxiv&utm_medium=paper&utm_campaign=covid-19. Potential pathogens were distinguished from commensal flora and contaminating microbial sequences from the environment and reagents by establishing a Z-score metric based on a background distribution derived from four non-templated “water-only” control libraries. Data were normalized to unique reads mapped per million input reads for each microbe at both species and genus levels. Taxa with Z-score less than 1, base pair alignment less than 50 base pairs, and reads per million (rpM) less than 10 were removed from analysis. Geneious Prime 2020.0.5 was used for separate de novo assembly and annotation of the draft SARS-CoV-2 genome. In subsequent runs to achieve higher coverage of the viral genome, two additional steps were taken: 1) the subsampling filter of the nonhost reads was removed and 2) targeted enrichment of the SARS-CoV-2 sequence using 73 tiling primers at a 10:1 molar ratio to random primers was adapted from viral genome recovery methods previously described.^6^

## Results

By February 1, the SARS-CoV-2 genome was correctly identified in the index COVID-19 patient sample using mNGS and globally accessible bioinformatics. An mNGS library was prepared, multiplexed, and sequenced. Notably, the sample generated 4,773,430 reads with 1,903,800 of those being non-host reads. The resulting non-host fastq files were processed using the IDseq platform which utilizes open-source components. When processed on IDseq pipeline version 3.18 with a February 2020 NCBI index, which included 54 SARS-CoV-2 sequences deposited by the international research community, sequences from the index patient aligned to the SARS-CoV-2 reference genome (Figure 1A, 582 reads, at 100% identity to the reference sequence; NCBI accession MN985325.1). Further analysis resulted in 33.2% coverage of the SARS-CoV-2 reference at an average depth of 2.2x (Figure 1B). Assembly and alignment on Geneious resulted in 50.5% coverage at 2.6x of the SARS-CoV-2 genome (NCBI accession MN908947.1). Re-sequencing the index case to a total sequencing depth of 8,786,916 single reads resulted in a greater number of SARS-CoV-2 alignments (819 reads) but had no impact on overall coverage – 32.3% coverage breadth at an average depth of 2.7x.

**Figure 1.**
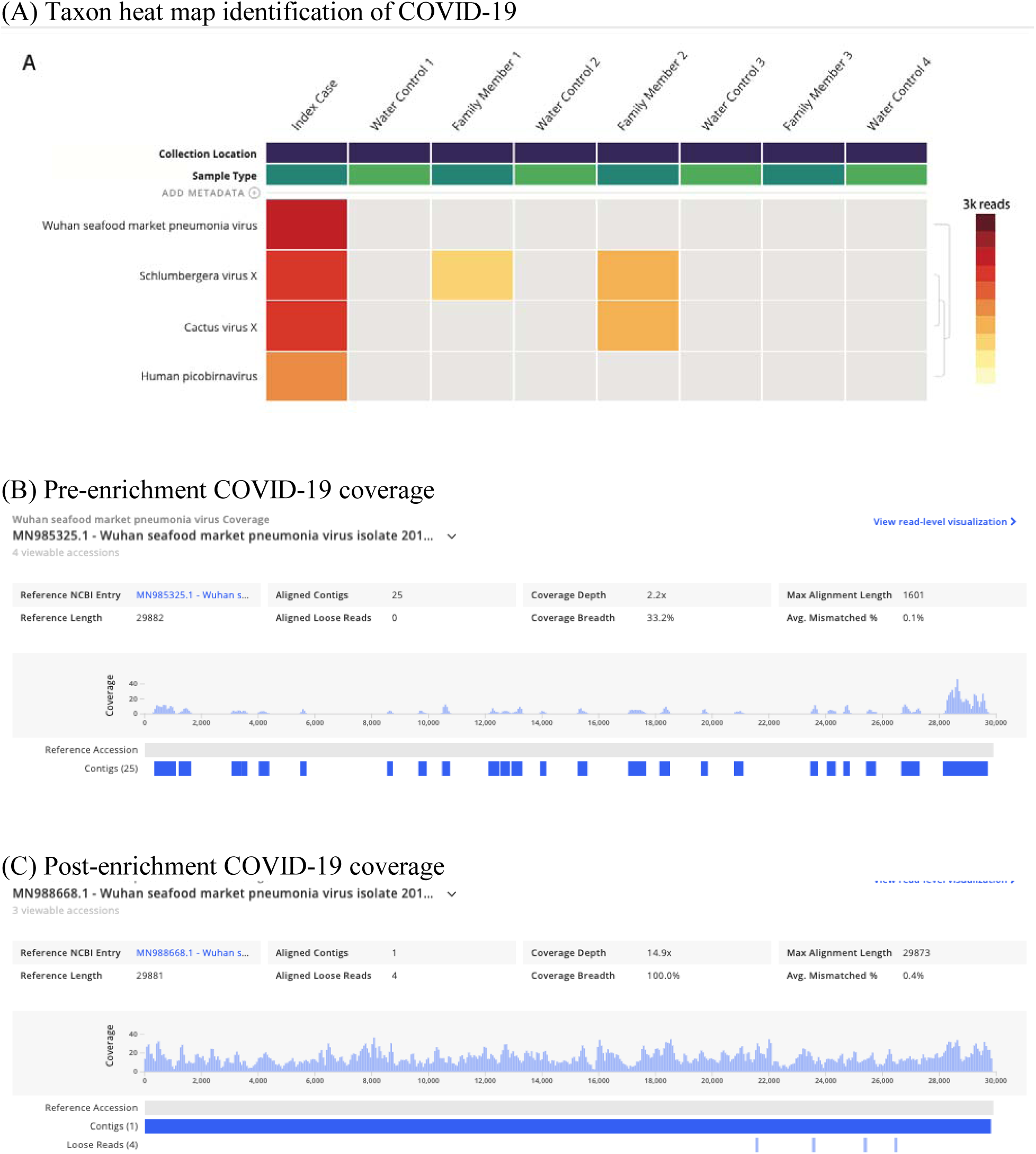
COVID19 viral genome identification and coverage. A) mNGS taxa heatmap results identifying COVID-19 virus; B) Coverage visualization map of COVID19 sequences; C) Coverage visualization map after target enrichment for COVID-19 sequences using spiked primer enrichment.

Using the target enrichment strategy, the overall fold enrichment was 6.4x. In terms of improving genome coverage, particularly at the 5’ end of the SARS-CoV-2 genome, the resulting 3,077 reads mapping to SARS-CoV-2 assembled into a single contig which provided full coverage of the reference genome at an average depth of 14.9x (Figure 1C)(NCBI accession MN988668.1).

All human SARS-CoV-2 genomes are very similar, including the SARS-CoV-2 genome from the Cambodian case as we would expect given that he transited directly from Wuhan. One SNP was noted at position 25,654 in ORF3a resulting in a valine-to-leucine substitution when compared to NCBI accession MN908947.1.

## Discussion

Here, we demonstrate that unbiased mNGS is a feasible and efficient means to detect pathogens in the midst of an outbreak of a novel virus. The index Cambodian SARS-CoV-2 sequence data uploaded to public databases represent one of the only sequences generated in a low-to-middle income country close to the outbreak epicenter. Despite major advances in mNGS technologies and significant decreases in costs associated with sequencing, preparation of sequencing libraries and sufficient bioinformatics capabilities for timely analysis still present a challenge in the developing world.^8^

As cases of COVID-19 spread globally in mid-January, initial real-time PCR protocols took into account that the genetic diversity of SARS-CoV-2 in humans and animals was not completely known. Non-specificity combined with variable sample quality can hinder ability to detect the pathogen in question. In this scenario, our metagenomics approach demonstrated a first-pass high coverage of the N gene, closely aligning to diagnostic protocols utilizing both the E and N genes for primary detection. This detection directly increased on-the-ground certainty in diagnostic sensitivity and specificity of available assays.^7^

mNGS in field settings proved critical in development of countermeasures for the 2014-2016 Ebola virus epidemic in West Africa, just as ongoing sequencing of COVID-19 cases from lesser developed areas of Southeast Asia will contribute to overall understanding of pathogen transmission, origin, and evolution.^9,10^ While COVID-19 cases are not limited to one remote region, as is often the case with viral hemorrhagic fever outbreaks, sequenced samples from all countries will be important for global disease containment. Instead of being limited to consensus Sanger sequencing, which may not detect high risk quasispecies, metagenomics is a powerful approach to detect variants of both novel and known species as epidemics evolve.^11^

The clear limitation in an mNGS approach is that low viral titers or high levels of host material demand greater read depth than may be available on instruments such as the iSeq100. To overcome this barrier, our follow-up steps included a target enrichment of SARS-CoV-2 while keeping the comprehensiveness of our mNGS pipeline intact.^6^ For an emerging threat, this strategy offers the flexibility to successfully recover the pathogen genome in question for subsequent phylogenetic analyses without compromising discovery. However, the other key factor in mNGS success is the accessibility to open-access, cloud-based metagenomics bioinformatics pipelines, such as IDseq which automates the process of separating host sequence characterizing the remaining non-host sequences.^12^ As recent as the Ebola epidemic, bioinformatics analyses were still mainly completed in the Global North.^10^ This newly available combination – more rugged, deployable sequencers plus user-friendly, globally accessible bioinformatics – represents an opportunity for responders in limited-resource settings; however, further proof-of-principle during outbreaks remains necessary.

Overall, agnostic or unbiased metagenomic sequencing capabilities in-country provide the ability to detect and respond to a variety of pathogens, even those that are unanticipated or unknown. Bridging of existing local and global resources for sequencing and analysis allows for better real-time surveillance locally, while also enabling better health pursuits overall, not just during outbreaks. The example described here serves as a call for continued training and infrastructure to support mNGS capacity in developing countries as bioinformatic tools proliferate and the cost of sequencing decreases.

## Acknowledgements

We would like to acknowledge the patient, Cambodian Ministry of Health and Cambodian Center for Disease Control, Rapid Response Team members, the Sihanoukville Provincial Health Department, and the doctors, nurses, and staff of Sihanoukville Hospital. We thank all of the technicians and staff at Institut Pasteur du Cambodge in the Virology Unit involved in this work especially Viseth Srey Horm, Kim Lay Chea, Phalla Y, Songha Tok, Sokhoun Yan, Sarath Sin, and Sonita Kol. We would like to thank Maira Phelps at Chan Zuckerberg Biohub for coordination of primers. The work of Dr. Manning is supported by the Division of Intramural Research at the National Institute of Allergy and Infectious Diseases at the National Institutes of Health and the Bill and Melinda Gates Foundation. The work of Dr. Karlsson is supported, in part, by, by the U.S. Department of Health and Human Services, Office of the Assistant Secretary for Preparedness and Response (Grant No. 1 IDSEP190051-01-00) and through internal funding at Institut Pasteur in Cambodia. The findings and conclusions in this report are those of the authors and do not necessarily represent the official views of the US Department of Health and Human Services.

